# Molecular basis for high affinity agonist binding in GPCRs

**DOI:** 10.1101/436212

**Authors:** Tony Warne, Patricia C. Edwards, Andrew S. Doré, Andrew G. W. Leslie, Christopher G. Tate

**Affiliations:** MRC Laboratory of Molecular Biology, Francis Crick Avenue, Cambridge CB2 0QH, UK; Heptares Therapeutics Ltd, BioPark, Broadwater Road, Welwyn Garden City, AL7 3AX, UK

## Abstract

A characteristic of GPCRs in the G protein-coupled state is that the affinity of the agonist often increases significantly, but the molecular basis for this is unclear. We have determined six active-state structures of the β_1_-adrenoceptor (β_1_AR) bound to conformation-specific nanobodies in the presence of agonists of varying efficacy. A direct comparison with structures of β_1_AR in inactive states bound to the identical ligands showed a 24-42% reduction in the volume of the orthosteric binding site. Potential hydrogen bonds were also shorter, and there was up to a 30% increase in the number of atomic contacts between the receptor and ligand. GPCRs are highly conserved, so these factors will likely be essential in increasing the affinity of a wide range of structurally distinct agonists.

**One Sentence Summary:** High affinity agonist binding to G protein-coupled GPCRs results from an increase in the number and strength of protein-ligand interactions.

## Main Text

GPCRs exist in an ensemble of conformations that can be selectively stabilized by the binding of a ligand and through interactions with signaling molecules such as G proteins (*1, 2*). The evidence for this comes from a wealth of pharmacological, biophysical and structural data. Pharmacology has characterized at least two distinct states of GPCRs, a conformation with high affinity for agonists when coupled to G proteins and a conformation with low affinity for agonists in the absence of G proteins (*1*), although a plethora of sub-states can also exist between these two extremes (*3, 4*). Fluorescence studies (*4, 5*) and ^19^F-NMR (*6, 7*) show that receptors in the absence of ligands can access many of these conformational states and that addition of an inverse agonist stabilizes a different state from an agonist or partial agonist. However, the fully active state can only be stabilized through coupling of a G protein or a G protein mimetic such as a conformation-specific nanobody (*8*). When GPCRs are stabilized in the fully active state, they typically show an increased affinity for the agonist, which can be as high as 100-fold, and has been observed in diverse receptors such as β_2_-adrenoceptors (*9*), the adenosine A_2A_ receptor (*10*), the muscarinic M2 receptor (*11*) and the μ-opioid receptor (*12*). In addition, the magnitude of the agonist affinity shift is dependent upon the structure of the ligand (*9*), although the molecular basis for this is unknown. Circumstantial evidence suggests that a decrease in the volume of the ligand binding pocket may be important during GPCR activation, although the exact nature of any effect is unclear because all previous comparisons have either been between different GPCRs (*13*) or between the same receptor but bound to ligands of different size and structures (*14*). Another proposal is that occlusion of the orthosteric biding site is the main cause for the increase in agonist affinity (*15*). There is thus currently no direct comparison between structures of a single receptor in the inactive state and the fully active state bound to the same ligand.

The majority of GPCR structures have been determined in an inactive state and these are highly conserved (*16*). A number of GPCR structures have also been determined in a fully active state coupled to either a G protein or a conformation-specific nanobody, revealing a conserved mechanism for GPCR activation (*17*). In addition, a number of structures have been determined of intermediate states (*18*), with agonists bound to receptors in the absence of a G protein or nanobody. In all these latter cases, the receptors are not in a fully active state and fall into two main classes, an active-like state or an inactive-like state. The active-like state is best characterised for the adenosine A_2A_ receptor (*19, 20*), where comparison with the G protein-coupled state (*10*) shows that full receptor activation is accompanied by the outward movement of H6 and the rotamer changes of Arg^3.50^ Tyr^5.58^ and Tyr^7.53^ (superscripts are the Ballesteros-Weinstein numbers (*21*)). Notably, there is no significant change in the structure of the orthosteric binding site (*10*). In contrast, structures of β-adrenoceptors bound to agonists (*22, 23*) are virtually identical to the inactive state bound to antagonists, except for a ~1 Å contraction of the orthosteric binding site and a rotamer change of Ser^5.46^. Previously we determined the structures of β_1_AR in an inactive state bound to agonists and partial agonists (*22, 24*) and this provides an ideal system for studying the molecular basis for agonists affinity shifts in GPCRs. We have therefore determined structures of β_1_AR in the active state coupled to a conformation-specific nanobody (either Nb80 or Nb6B9) used previously to crystallise β_2_AR in an active state coupled to agonists (*13, 25*).

Six crystal structures with overall resolutions between 2.9 Å - 3.2 Å (Table S1) were determined of β_1_AR bound to either Nb80 or Nb6B9, and the overall structures were all virtually identical (Fig. 1; 0.2-0.3 Å RMSD for Cα atoms). Structures were determined bound to full agonists (isoprenaline, formoterol), partial agonists (salbutamol, dobutamine, xamoterol) and a weak partial agonist formerly described as an antagonist (cyanopindolol). All the agonists and partial agonists showed an increase in affinity when β_1_AR was coupled to the engineered G protein mini-G_s_, whereas cyanopindolol bound with similar high affinity in both the presence and absence of mini-G_s_ (Fig. 1). The inability of mini-G_s_ to increase the affinity of cyanopindolol was not a consequence of the oxymethylene spacer between the ethanolamine backbone and ligand head group that prevents contraction of the ligand-binding pocket in β_1_AR antagonists, because this is also found in the partial agonist xamoterol. The overall structure of the β_1_AR-nanobody complexes bound to either agonists or partial agonists is virtually identical to that of the agonist-bound Nb6B9-β_2_AR complex (0.5 Å RMSD of 1601 Cα atoms) and the overall conformational changes compared to the inactive state are consequently very similar (Fig. 2). These are characterised by an outward movement of the cytoplasmic ends of H5 and H6 and an inward movement of H7. In contrast, the extracellular ends of H6 and H7 move inwards, with sideways movements of the extracellular ends of H1 and H2 and an inwards movement of ECL2. These changes result in the partial occlusion of the orthosteric binding pocket (Fig. 2), which is consistent with observations on nanobody-bound β_2_AR (*13, 25*).

**Fig. 1.**
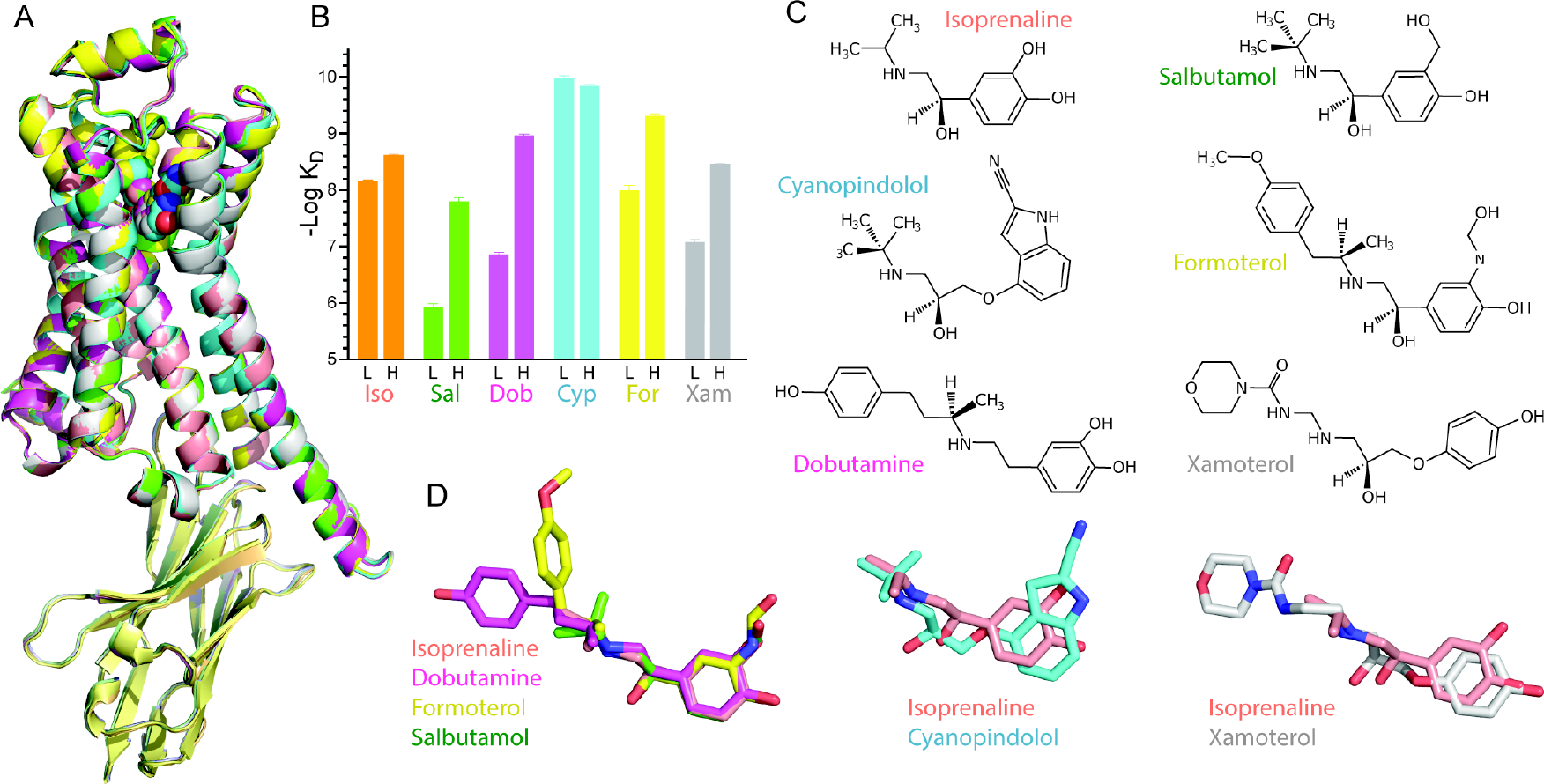
Structure of the active state of agonist-bound β_1_AR-nanobody complex. (A) Superposition of six structures of β_1_AR complexes bound to ligands shown in (C). (B) Affinities of β_1_AR in the low affinity state, L, and high affinity state coupled to mini-G_s_, H, for the ligands co-crystallised with the receptor: Iso, isoprenaline (orange); Sal, salbutamol (green); Dob, dobutamine (purple); Cyp, cyanopindolol (cyan); For, formoterol (yellow); Xam, Xamoterol (grey). Data are in Tables S1 and S2 and representative graphs of affinity shifts are in Fig. S4. Results are the mean of 2-4 experiments performed in duplicate with error bars representing the SEM. (C) Structures of the ligands co-crystallised in the β_1_AR complexes. (D) Disposition of the ligands after superposition of the receptors, using the same colour coding as in (B).

**Fig. 2.**
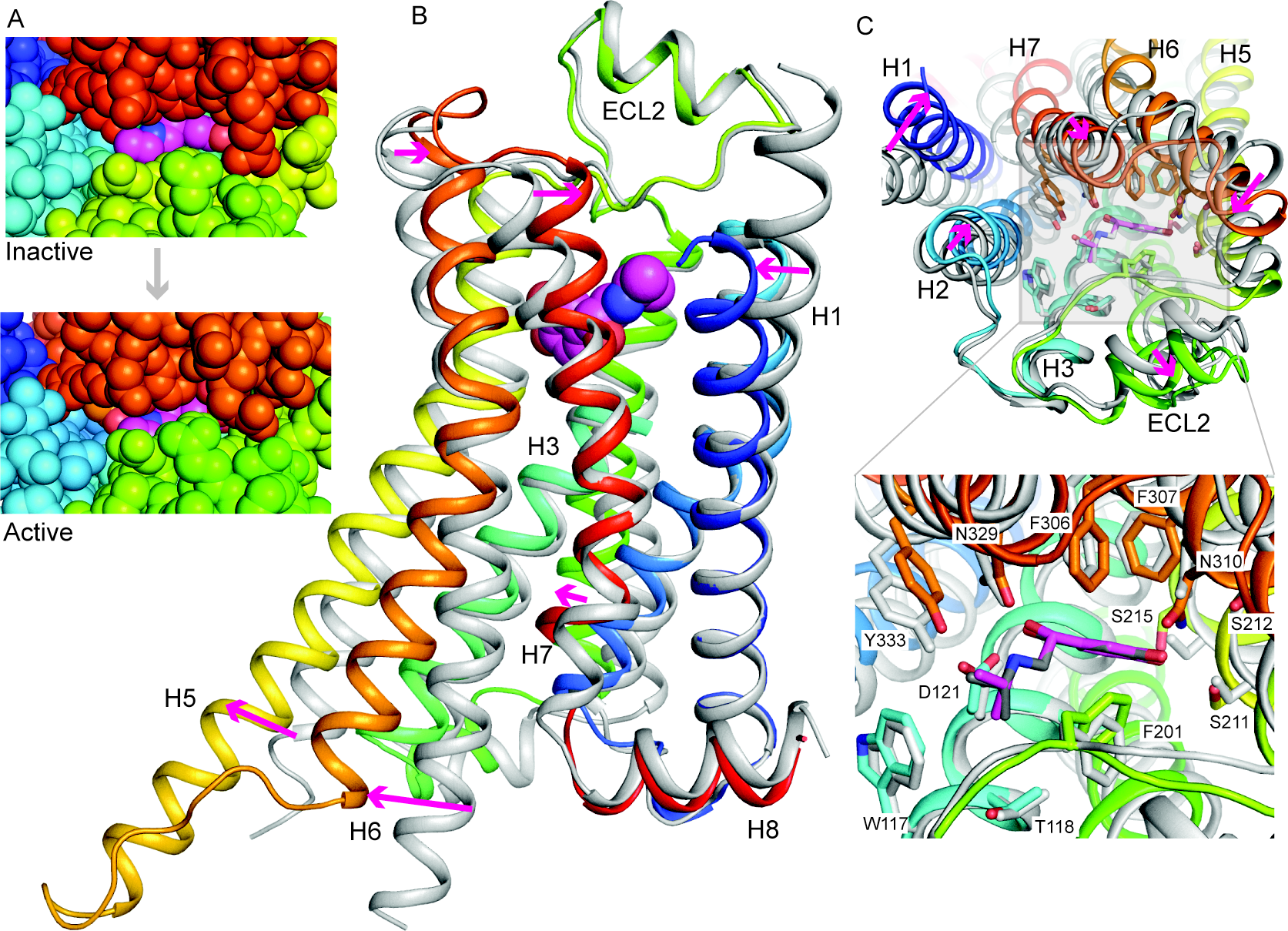
Conformational changes in isoprenaline-bound β_1_AR. (A) View of the orthosteric binding site from the extracellular surface with atoms shown as space filling models: isoprenaline (magenta, carbon atoms); β_1_AR: H1, dark blue; H2, light blue; H5, yellow; ECL2, green; ECL3 and parts of H6 and H7, brown. (B-C), Superposition of isoprenaline-bound β_1_AR in the inactive state (grey, PDB ID 2Y03) with isoprenaline-bound β_1_AR in the active state (rainbow colouration; Nb6B9 has been removed for clarity). Arrows (magenta) indicate the transitions from the inactive to active state. Alignment was performed based on the isoprenaline molecules using PyMol (magenta, isoprenaline bound to active state β_1_AR). (B) β_1_AR viewed parallel to the membrane plane. (C) β_1_AR viewed from the extracellular surface with the inset showing residues lining the orthosteric binding site.

Detailed comparisons were made between the inactive state structures of β_1_AR bound to either isoprenaline, salbutamol, dobutamine or cyanopindolol with the respective active state structures (Figs. 3 and 4). In all cases, there was a decrease in the volume of the orthosteric binding site that varied depending on the ligand (Fig. S1). The largest decrease was observed for the full agonist isoprenaline and the smallest decrease was observed for the weak partial agonist cyanopindolol (volume reductions: isoprenaline, 42%, dobutamine, 30%; salbutamol, 30%; cyanopindolol, 24%). The decrease in the volume of the orthosteric binding site when isoprenaline was bound was due primarily to the inward movement of the extracellular ends of H6 and H7, an inward movement and an increase in the H5 bulge at Ser215^5.46^ and the reorientation of residues Phe201^ECL2^ and Phe325^7.35^. The magnitude of these changes correlated with efficacy: thus the structure with cyanopindolol bound showed little change at the extracellular side of H7 (0.9 Å shift when cyanopindolol was bound compared to 3.7 Å when isoprenaline was bound; measurement at the Cα of Asp322^7.32^) and less change in the bulge of H5 (2.0 Å shift when cyanopindolol was bound compared to 3.4 Å when isoprenaline was bound; measurement at Cα of Ser215^5.46^). The pincer-like movement of Phe201^ECL2^ and Phe325^7.35^ towards the ligand has the largest effect on reducing the volume of the orthosteric binding pocket, with the maximal shift observed in the isoprenaline structure of 3.1 Å for Phe201^ECL2^ and 2.5 Å for Phe325^7.35^ (measured at the CZ atom of the side chain). The movement of Phe201^ECL2^ appeared to correlate with the ligand structure because in all cases it formed van der Waals contacts with the ligand. In contrast, Phe325^7.35^ was not within van der Waals contact with any of the four ligands and moved as a consequence of the inward tilt of H7.

**Figure 3.**
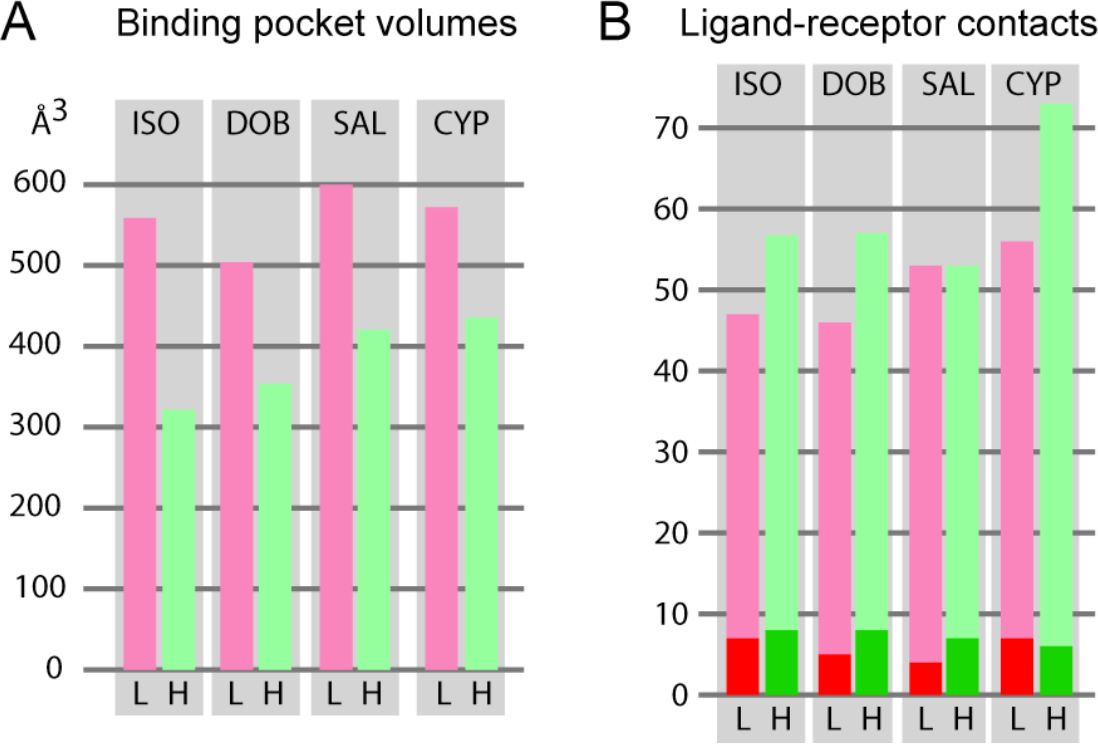
Analyses of the orthosteric binding site in active state structures of β_1_AR. (A) Volume differences of the orthosteric binding site in the low-affinity inactive state (L, pink bars) compared to the high-affinity active state (H, green bars). (B) Number of atomic contacts between the respective ligands and β_1_AR in the low-affinity inactive state (L, pink bars) compared to the high-affinity active state (H, green bars). The dark shades represent the number of polar interactions Ligand abbreviations are shown in Fig. 1.

The reduction in the volume of the orthosteric binding pocket correlated with an overall reduction in the average distance between atoms in the ligand and receptor by 0.1-0.3 Å. Although amino acid residues in H3, H5, H6, H7 and ECL2 (and H2 for dobutamine) were all involved in contributing to ligand-receptor contacts, the biggest decreases in contact distance were ligand dependent (Fig 4). The greatest decreases in ligand-receptor distances were observed between Asp121^3.32^ and salbutamol, Gly98^2.61^ and dobutamine, and Val125^3.36^ and cyanopindol, where the residues were all greater than 1 Å closer to the ligand in the active state. However, the changes around H5 and H6 may lead to greater changes in affinity as these involved the strengthening of hydrogen bonds. For example, Asn310^6.55^ was predicted to make a weak hydrogen bond to the para-hydroxyl group of isoprenaline (3.5 Å between donor and acceptor) in the inactive state, which changes to 2.8 Å in the active state. In contrast, hydrogen bonds formed by Ser211^5.42^ and Ser215^5.46^ do not alter length significantly. This differed to observations for dobutamine and salbutamol where the hydrogen bond to Ser215^5.46^ was 0.8 Å shorter for both ligands and the hydrogen bond to Ser211^5.42^ also shortened by 0.7 Å to salbutamol, but remained unchanged to dobutamine. Most of the observed differences are due to the contraction of the binding pocket, but the significant shortening of the hydrogen bond between Ser211^5.42^ and salbutamol is due to a rotamer change.

**Figure 4.**
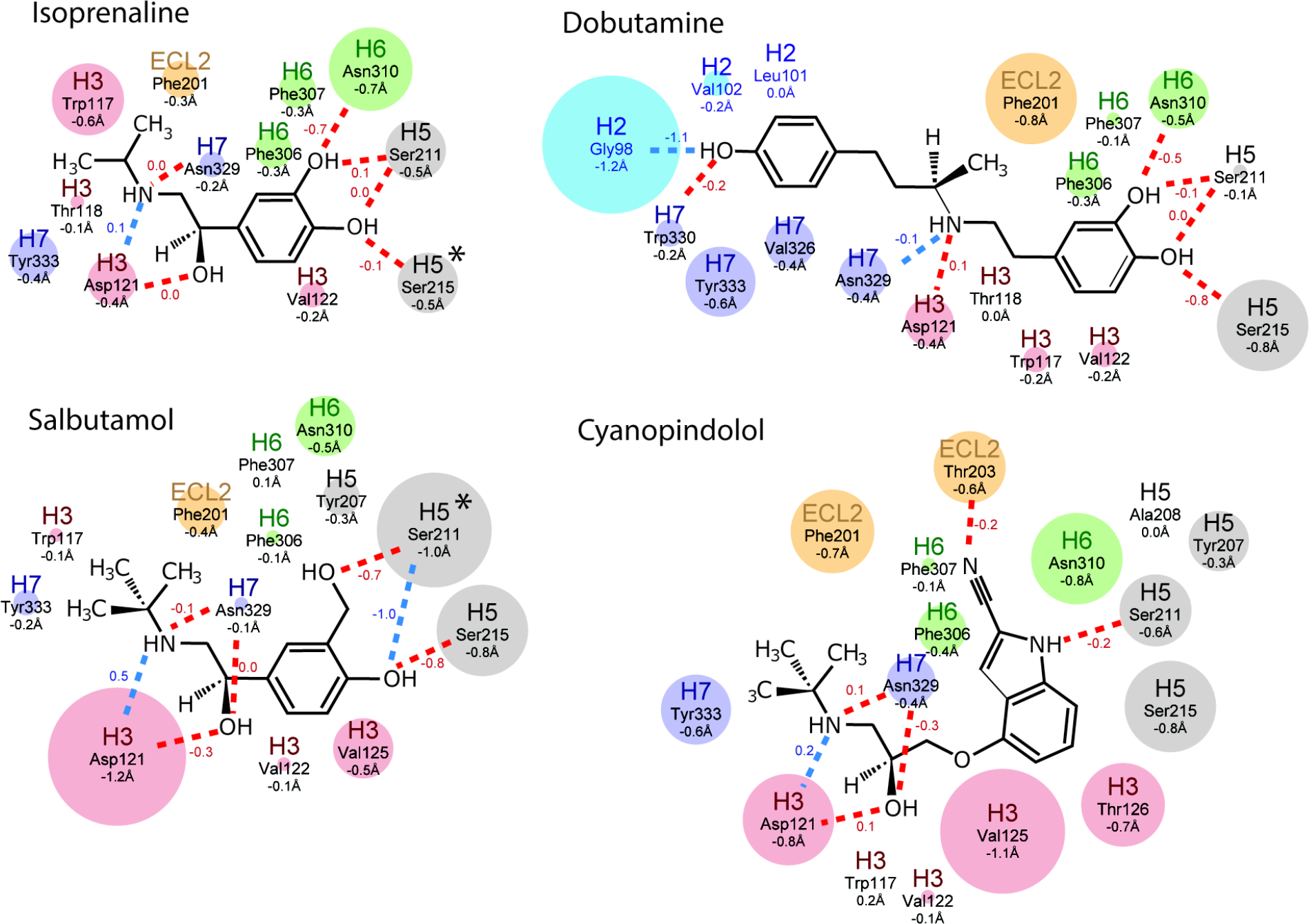
Changes in receptor-ligand contact distances. The maximal changes in contact distances between ligands and atoms in β_1_AR from the inactive to active states are depicted. Amino acid side chains making contact to the ligands are indicated and coloured according to where they are in β_1_AR (blue, H2; red, H3; orange, ECL2; grey, H5; green, H6; purple, H7) with the diameter of the circle representing the magnitude of the distance change (shown as numbers below the amino acid residue). Numbers next to the lines indicate the change in length of polar contacts (blue dashed lines) and hydrogen bonds (red dashed lines; determined using HBPLUS). Negative numbers imply a decrease in distance between the ligand and receptor fin the transition from the inactive state to the active state. An asterisk indicates a significant rotamer change between in the inactive and active states. For the disposition of additional contacts made by each side chain, see Fig. S2.

There were also notable increases in the number of ligand-receptor contacts made by Phe201^ECL2^ to cyanopindol and Asp121^3.32^ to salbutamol that were not observed at the respective positions with the other ligands (Fig. S2). Thus, although all the ligand binding pockets contracted upon receptor activation, the changes in ligand-receptor contacts were not conserved, despite the similarity in chemotypes amongst the four ligands studied. In addition, there was no clear correlation between the number and type of ligand-receptor interactions present and either the magnitude of ligand affinity increase on receptor activation or the decrease in volume of the orthosteric binding site. It was particularly notable that cyanopindolol bound to β_1_AR with similar affinity in both the presence and absence of a coupled G protein despite the contraction of the binding pocket and increase in receptor ligand contacts upon activation. This may be a consequence of constraints on the possible conformation change imposed by the rigidity of cyanopindol that prevents the full contraction of the ligand binding pocket by preventing the movement of H7 and the bulge in H5 that are observed in the other structures (Fig. S1).

Previous structural studies on β_1_AR suggested that the mode of ligand interaction with Ser215^5.46^ in an inactive state correlates with efficacy by affecting the likelihood of transitions to activated states (*22, 26*). The active state structures determined here suggest that the ability of ligands to stabilize the activated state must also be taken into account when considering ligand efficacy. For example, the weak partial agonists cyanopindolol and xamoterol do not allow the full contraction of the ligand binding pocket observed with isoprenaline or formoterol. In contrast, the partial agonists salbutamol and dobutamine show similarly contracted binding pockets to isoprenaline and formoterol, but do not engage Ser215^5.46^ in the inactive state like full agonists (*22*).

The role of the partial occlusion of the orthosteric binding site upon activation of β_1_AR was tested by mutagenesis inspired from the active state structure of β_2_AR (*13, 25*). In β_2_AR, it was proposed that the occlusion of the binding site was a significant factor in increasing agonist affinity upon G protein coupling (*15*). In particular, Tyr308^7.35^ was within van der Waals distance of Phe193^ECL2^ on the opposite side of the entrance to the orthosteric binding pocket and had a major effect on decreasing the rates of association and dissociation of ligands in the active state compared to the inactive state. The β_2_AR residues Phe193^ECL2^/Tyr308^7.35^ are equivalent to Phe201^ECL2^/Phe325^7.35^ in β_1_AR where they are not in van der Waals contact and therefore do not occlude significantly the entrance to the binding pocket as observed in β_2_AR (Fig. 5). Thus the mutation F325Y^7.35^ in β_1_AR was predicted to occlude the entrance to the orthosteric binding pocket and decrease the rate of ligand association, and conversely, F325A^7.35^ was predicted to make the entrance wider and increase the rate of ligand association. When the initial rate of ^3^H-dihydroalprenolol (^3^H-DHA) association was measured (Fig. 5), β_1_AR(F325A) had the same rate as β_1_AR, but β_1_AR(F325Y) had a considerable slower rate of association. However, the affinities (Fig 5) of epinephrine and isoprenaline for the high affinity state of β_1_AR and β_1_AR(F325Y) were identical (epinephrine, 3 nM and 3.3 nM, respectively; isoprenaline, 2.4 nM and 2.0 nM, respectively) and there was a small difference with norepinephrine (1.2 nM and 3.2 nM, respectively). Comparisons of affinities (Fig 5) for the inactive states of β_1_AR and β_1_AR(F325Y) showed a large decrease in affinities for norepinephrine (5.8 fold), epinephrine (10.5-fold) and isoprenaline (6.5-fold), which implied that the greater agonist affinity shift observed in β_1_AR(F325Y) compared to β_1_AR was due to destabilisation of the inactive state and not stabilisation of the active state. This suggested that partial occlusion of the ligand binding pocket in β_1_AR(F325Y) during formation of the active state played little role in the increase of agonist affinity on G protein coupling.

**Figure 5.**
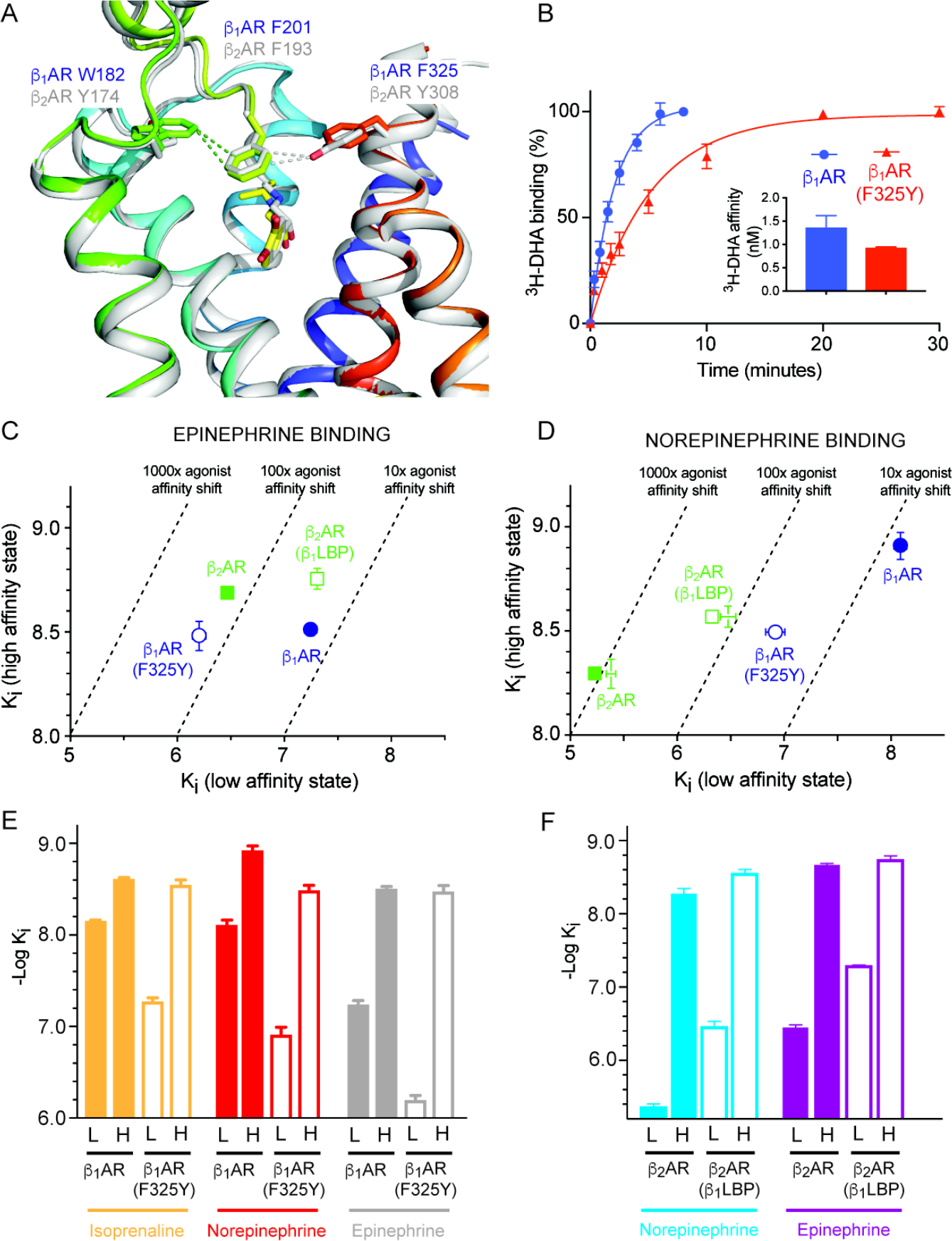
Comparisons between β_1_AR and β_2_AR. (A) The active state structures of β_1_AR (rainbow colouration) and β_2_AR (grey, PDB ID 4LDO) were aligned and positions of key residues in the extracellular surface depicted. Ligands are shown as sticks; isoprenaline, yellow; adrenaline, grey. (B) Rate of association of the radioligand ^3^H-DHA on to β_1_AR (blue circles) and β_1_AR(F325Y) (red triangles) and β_1_AR(F325A) (orange circles). The inset shows the affinities of ^3^H-DHA for β_1_AR (blue bars) and β_1_AR(F325Y) (red bars) and β_1_AR(F325A) (orange bars). (C-D) Comparison of the affinity for agonist binding in the high affinity state and the affinity for agonist binding in the low affinity state; β_1_AR, blue filled circles; β_1_AR(F325Y), blue open circles; β_2_AR green filled squares; β_2_AR(β_1_LBP), green open squares (Tables S2 and S3; Figure S4). (E-F) Affinities of β_1_AR, β_2_AR and their respective mutants in the low affinity state, L, and high affinity state coupled to mini-G_s_, H. All data are in Tables S1 and S2 and representative graphs of affinity shifts are in Figure S4. Results are the mean of 2-7 experiments performed in duplicate with error bars representing the SEM.

The destabilising effect of the F325Y mutation in β_1_AR on the agonist-bound inactive state suggested that converting the extracellular surface of β_2_AR to make it similar to β_1_AR would increase the affinity of the inactive state and leave the affinity of the G protein-coupled activated state approximately unchanged. The β_2_AR mutant constructed, β_2_AR(β_1_LBP), did indeed show these characteristics (Fig 5; see Methods for the rationale of the four mutations used: Y174W^ECL2^, H296N^6.58^, K305D^7.32^ and Y308F^7.35^). In addition, the accessibility of the β_1_AR orthosteric binding pocket in the G protein-coupled state to ^125^I-cyanopindolol was greater than that observed for β_2_AR (Fig. S3). The four mutations in β_2_AR(β_1_LBP) converted the behaviour of β_2_AR to that of β_1_AR. Adding the converse residues from β_2_AR into β_1_AR, to make the mutant β_1_AR(β_2_LBP), converted the accessibility of the orthosteric binding site in β_1_AR to that of β_2_AR (Fig. S3).

The multiple structures of β_1_AR in the activated state bound to ligands of different efficacy defined major contributors towards the increase in agonist affinity upon G protein coupling. The orthosteric binding pocket decreased in volume, regardless of the efficacy of the ligand bound. In addition, there were more ligand-receptor contacts and/or interactions of greater strength due to shortening of the contacts involved. Partial occlusion of the entrance to the receptor binding pocket was excluded as a potential factor that affected the change in agonist affinity on G protein coupling, although it was clearly a factor in changing the kinetics of ligand association. A purely steric effect would not be expected to affect ligand affinity, given that this is the ratio between the association and dissociation constants, but it is apparent from mutagenesis data that mutations of key residues in the entrance to the orthosteric binding site preferentially affected the agonist affinity of the inactive state, and hence altered the agonist affinity shift upon G protein coupling. A key finding of this work was that the increase in agonist affinity upon G protein coupling arose from changes at different amino acid residues in different areas of the orthosteric binding pocket. This has implications for drug development as structure-based drug design usually considers only a single state, despite the fact that any drug will actually experience multiple different states of a GPCR. In the absence of crystal structures of multiple states of most receptors with multiple different ligands, the insights into how ligands bind to both the inactive and active states of β_1_AR will help in developing tools for engineering efficacy into ligands at an early stage in drug development. As GPCRs are highly conserved, these conclusions are likely to be applicable to many different receptors.

## Acknowledgments

We thank the beamline staff at the European Synchrotron Radiation Facility (beamlines ID23-2, ID30-A3, ID29, ID30B and MASSIF-1) and at Diamond Light Source (beamline I24).

## Funding

This work was supported by core funding from the Medical Research Council [MRC U105197215 and U105184325] and a grant from the ERC (EMPSI 339995);

## Author contributions

T.W. performed receptor and nanobody expression, purification, crystallization, cryo-freezing of the crystals, data collection, data processing and structure refinement. T.W. also performed the pharmacological analyses. P.C.E. purified mini-G_s_ and A.D.S assisted with structure solution. A.G.W.L. was involved in data processing and structure solution, refinement and analysis. Manuscript preparation was performed by T.W., A.G.W.L. and C.G.T. The overall project management was by C.G.T.;

## Competing interests

C.G.T. is a shareholder, consultant and member of the Scientific Advisory Board of Heptares Therapeutics, who also partly funded this work;

## Data and materials availability

The co-ordinates and structure factors for all the structures determined have been deposited at the PDB with the following accession codes (ligand co-crystallised in parentheses): 6H7J (isoprenaline), 6H7K (formoterol), 6H7L (dobutamine), 6H7M (salbutamol), 6H7N (xamoterol), 6H7O, cyanopindolol.

## Supplementary Materials

### Materials and Methods

#### Cloning, expression and purification of β_1_AR

The turkey (*Meleagris gallopavo*) β_1_AR construct trx-β_1_AR (*22*) used for crystallization of the β_1_AR-nanobody complexes was based on β44-m23, with the same truncations and deletions, but only four thermostabilizing mutations, R68S^1.59^, M90V^2.53^, F327A^7.37^ and F338M^7.48^. The mutations Y227A^5.58^ and A282L^6.27^ on H5 and H6 were removed, because the reversion of these two mutations was sufficient to enable full activation and high affinity agonist binding in the presence of G proteins and nanobody Nb80 (*27*). A thioredoxin (*E. coli* trxA, with mutations C32S & C35S) fusion was attached via the linker EAAAK at the N-terminus of β_1_AR. The construct was cloned into the baculovirus transfer vector pAcGP67B (BD Biosciences) and the recombinant baculovirus was generated by co-transfection of insect cells with BacPAK6 linearized baculovirus DNA (Oxford Expression Technologies Ltd). Plaque purified virus was used to express receptors in High Five cells (ThermoFisher Scientific) grown in ESF921 (Expression Systems) supplemented with 5% heat-inactivated foetal bovine serum (Sigma) as described previously (*28*).

The membrane fraction was prepared, and the receptor was solubilized in 1.5% decylmaltoside (DM, Generon) and further purified in 0.1% DM by Ni^2+^-affinity chromatography and alprenolol sepharose chromatography, with elution from the alprenolol sepharose ligand affinity column as described previously (*22, 28, 29*) with 100 μM of the appropriate ligand for complex formation, concentrated to 15-25 mg/ml and either used directly for the formation of complexes, or frozen for later use.

#### Expression and purification of nanobodies Nb80 and Nb6B9

Synthetic genes (Integrated DNA Technologies) for Nb80 (*30*) and Nb6B9 (*13*) were cloned into plasmid pET-26b(+) (Novagen) with a N-terminal His_6_ tag followed by a thrombin protease cleavage site. Expression in *E. coli* strain BL21(DE3)RIL (Agilent Technologies) and purification from the periplasmic fraction were as described elsewhere (*13*), but with the addition of a final thrombin (Sigma) protease cleavage step to remove the His_6_ tag before concentration to 40 mg/ml.

#### Formation of agonist-bound trx-β_1_AR-nanobody complexes and purification with detergent exchange by size exclusion chromatography (SEC)

Trx-β_1_AR (1.0-2.0 mg) was mixed with 1.5-fold molar excess nanobody (0.4-0.8 mg) with the addition of cholesteryl hemisuccinate (Sigma) to 0.1 mg/ml in a final volume of 150 μL. For the formation of activated complexes with receptor purified in full agonists (isoprenaline and formoterol), Nb80 was used and incubation was for 2 hours at room temperature. For the formation of complexes with receptor purified in partial agonists (salbutamol, dobutamine, xamoterol and cyanopindolol), trx-β_1_AR was mixed with Nb6B9 and incubated overnight at room temperature. After incubation, size exclusion chromatography (SEC) was performed to separate receptor-nanobody complexes from excess nanobody and to exchange the detergent from DM to HEGA-10 for crystallization by vapour diffusion. A Superdex Increase 200 10/300GL column (GE) was used at 4°C, the column was equilibrated with SEC buffer (10 mM Tris-HCl pH 7.4, 100 mM NaCl, 0.1 mM EDTA, 0.35% HEGA-10 [Anatrace]) supplemented with 10 μM of the appropriate agonist ligand. Samples containing complex were mixed with 200 μL SEC buffer and centrifuged (14,000 × *g*, 5 minutes) immediately prior to SEC (flow rate 0.2ml/minute), with a run time of one hour which was sufficient for a near-complete detergent exchange as indicated by quantitation of residual glycosidic detergent (*31*). Peak fractions corresponding to complex were concentrated to 15 mg/ml for crystallization by vapour diffusion using Amicon Ultra-4 50 kDa centrifugal filter units (EMD-millipore).

#### Crystallization of receptor-nanobody complexes, data collection, processing and refinement

Crystals were grown in 150 + 150 nL sitting drops by vapour diffusion at 18°C against reservoir solutions containing 0.1 M Hepes-NaOH pH 7.5 and 21-24% PEG1500; the yield of crystals was increased by addition of HEGA-10 to 0.5-0.6% prior to setting up the drops. Crystals usually appeared within 2 hours and grew to full size (up to 200 μm in length) within 48 hours. Crystallization plates were equilibrated to 4°C for at least 24 hours before cryo-cooling. Crystals were picked in LithoLoops (Molecular Dimensions Ltd) and transferred to 0.1 M Hepes-NaOH pH 7.5, 25% PEG1500 containing 5% glycerol for 2 seconds before plunging into liquid nitrogen.

Diffraction data for trx-β_1_AR-nanobody complex crystals were collected at ESRF, Grenoble using beamlines id23-2, id30-a3, id29, id30b & MASSIF-1. Helical collection strategies were used to collect complete data sets while translating between two points in order to minimize radiation damage, except for trx-β_1_AR-Nb80-formoterol where X-ray diffraction data were collected from a single point on the crystal by the autonomous ESRF beamline MASSIF-1 (*32*) using automatic protocols for the location and optimal centering of crystals (*33*). The beam diameter was selected automatically to match the crystal volume of highest homogeneous quality and was therefore collimated to 30 μm, and strategy calculations accounted for flux and crystal volume in the parameter prediction for complete data sets(*34*). Although the thioredoxin fusion was not well resolved in the structures, it was important for ease of data collection as it resulted in crystals with an orthorhombic space group and not monoclinic as is usual when β_1_AR is crystallized in Hega-10 (*22*). Diffraction data were processed using MOSFLM (*35*) and AIMLESS (*36*), structures were solved using PHASER (*37*) with use of the crystal structures of the active state β_2_AR stabilized with nanobody Nb80(*30*) and wild-type thioredoxin (PDBs 3P0G and 2H6X) as search models. Diffraction was anisotropic, so merged data were analyzed and subjected to anisotropic truncation using the UCLA Diffraction Anisotropy Server (http://services.mbi.ucla.edu/anisoscale/) and STARANISO (http://staraniso.globalphasing.org). Model refinement and rebuilding were carried out with REFMAC5 (*38*) and COOT (*39*).

#### Expression and purification of mini-G_s_

Mini-G_s_ (construct 393) was expressed in *E. coli* strain BL21(DE3)RIL and purified by Ni^2+^-affinity chromatography, followed by cleavage of the histidine tag using TEV protease and negative purification on Ni^2+^-NTA to remove TEV and undigested mini-G_s_; SEC was then used to remove aggregated protein as described elsewhere (*40*). Purified mini-G_s_ was concentrated to give a final concentration to 100 mg/ml in 10 mM HEPES, pH 7.5, 100 mM NaCl, 10% v/v glycerol, 1 mM MgCl_2,_ 1 μM GDP and 0.1 mM TCEP.

#### Preparation of activated trx-β_1_AR-mini-Gs complexes

For the comparison of complex formation with mini-G_s_ in the presence of either full or weak partial partial agonist, 150 μM trx-β_1_AR was incubated overnight at 4°C with 200 μM mini-G_s_ in a final volume of 200 μL SEC buffer (10 mM Tris-HCl pH7.4, 100 mM NaCl, 1 mM MgCl_2_, 0.1% DM) containing 0.75 mM ligand (either isoprenaline or cyanopindolol). A further 1 hour incubation followed addition of 0.1 unit apyrase (Sigma), after which the sample was centrifuged (14,000 ×*g*, 5 minutes) before SEC using a Superdex Increase 200 10/300GL column (GE) at 4°C. The column was run at 1 ml/minute in SEC buffer with the addition of 10 μM ligand, and 0.8ml fractions were collected for analysis by SDS-PAGE. The results of these experiments are shown in Fig. S5, and indicate that in the presence of weak partial agonist, the trx-β_1_AR-mini-G_s_ complex is unstable, and therefore G protein mimetic nanobodies Nb80 or Nb6B9 were used in this study to prepare crystals with a range of ligands with differing pharmacological profiles.

#### Radioligand binding studies on βARs and mutants

Wild type turkey β_1_AR, human β_1_AR and human β_2_AR, and mutants of these receptors, were all expressed using recombinant baculoviruses in insect cells for radioligand binding studies. Amino acid residues close to the ligand binding pocket (LBP) which differ between β_1_AR and β_2_AR were mutated to compare some of the pharmacological characteristics of the different receptor subtypes. The residues selected for mutation were Trp182^ECL2^, Asn313^6.58^, Asp322^7.32^ and Phe325^7.35^ (β_1_AR), equivalent to Tyr174^ECL2^, His296^6.58^, Lys305^7.32^ and Tyr308^7.35^ (β_2_AR). The first residues, Trp182^ECL2^/Tyr174^ECL2^ (β_1_AR/β_2_AR) was chosen because they are involved in differing modes of interaction with Phe201^ECL2^/Phe193^ECL2^ that were observed in comparisons of crystal structures, as well as possible involvement in a secondary affinity state observed in β_1_AR but not β_2_AR(*41*). The latter three pairs of residues were chosen because His296^6.58^, Lys305^7.32^ and Tyr308^7.35^ have all been suggested to contribute to high affinity binding of agonist to β_2_AR in the presence of G protein (*13, 15*). For further explanation see Fig S6.

The initial turkey β_1_AR construct was based on the β44-m23 construct (*22*), but without any of the stabilizing mutations. Two variants of β_1_AR were prepared with mutations of amino acids in the ligand binding pocket (LBP) that were intended to make the β_1_AR similar to the β_2_AR. These were β_1_AR(F325Y) (mutation F325Y^7.35^) and β_1_AR(β_2_LBP) that contained following mutations: W182Y^ECL2^, N313H^6.58^, D322K^7.32^, F325Y^7.35^. The human β_2_AR was mutated to generate the construct β_2_AR(β_1_LBP) that contained the mutations Y174W^ECL2^, H296N^6.58^, K305E^7.32^ and Y308F^7.35^. Mutants were constructed in the baculovirus transfer vectors pBacPAK8 (Clontech) for β_1_AR and pAcGP67B (BD Biosciences) for β_2_AR(β_1_LBP) by using Quikchange protocols (Stratagene) with KOD polymerase (EMD Millipore), and were expressed in insect cells after co-transfection with linearized baculovirus as previously described. Crude insect cell membrane fractions were prepared by resuspending cell pellets from 1 ml culture volume in 1 ml of assay buffer (20 mM Hepes-NaOH pH7.5, 50 mM NaCl, 2.5 mM MgCl_2_, 0.1% BSA) to give final concentrations of 1-3 × 10^6^ cells/ml. Cells were sheared by 10 passages through a bent 26G needle and cell debris was removed by centrifugation (1500 x*g*, 2 min) and the supernatants were diluted in assay buffer for radioligand binding studies.

#### Saturation binding assay to determine affinities for [^3^H]-dihydroalprenolol and [^125^I]-cyanopindolol

Saturation binding assays were performed on all constructs to determine appropriate apparent K_D_ values for [^3^H]-dihydroalprenolol (DHA) to use in competition binding assays. Insect cell membranes containing βAR constructs were diluted in 20 mM Hepes-NaOH pH 7.5, 50 mM NaCl, 2.5 mM MgCl_2_, 1 mg/ml BSA. The sample was aliquoted and dilutions of [^3^H]-DHA (Perkin Elmer) were added to give final concentrations in the range of 0-20 nM (β_1_AR constructs) and 0-2.5nM (β_2_AR constructs) in a final volume of 220 μL, with 10 determinations in duplicate per binding curve. Non-specific binding was determined by addition of alprenolol to negative controls (1 mM final concentration). Samples were incubated at 20°C for 2 h, before filtering 100 μL duplicate aliquots through 96-well Multiscreen HTS GF/B glass fibre filter plates (Merck Millipore) pre-soaked in 0.1% w/v polyethyleneimine to separate bound from unbound [^3^H]-DHA. Filters were then washed three times with 200 μL volumes ice-cold assay buffer (20 mM Hepes-NaOH pH 7.5, 50 mM NaCl, 2.5 mM MgCl_2_). Filters were dried, punched into scintillation vials and 4 ml Ultima Gold scintillant (Perkin Elmer) were added. Radioligand binding was quantified by scintillation counting using a Tri-Carb Liquid Scintillation Analyser (Perkin Elmer) and apparent K_D_ values were determined using GraphPad Prism version 7.0b (GraphPad Software, San Diego, CA). Apparent K_D_ values were determined for all constructs, also in the presence of 25 μM mini-G_s_ (R393), in which case apyrase (0.1 U/ml final concentration) was included. All K_D_ values obtained are given in Tables S2 and S3, and are mean values obtained from at least two experiments performed in duplicate. Mean K_D_ values for [^125^I]-cyanopindolol (Cyp) were also determined for β_1_AR using a concentration range of 0-1000 pM [^125^I]-Cyp, with non-specific binding determined by addition of cyanopindolol to negative controls (0.1 mM final concentration).

#### Competition binding assays

Insect cell membranes containing βAR were resuspended in assay buffer (20 mM HEPES pH 7.5, 50 mM NaCl, 2.5 mM MgCl_2_, 0.1% BSA, with inclusion of 0.1 mM ascorbate for ligands containing catechols). The sample was aliquoted and mini-G_s_ construct R393 (25 μM final concentration, for determination of high affinity states), agonist (8 points with final concentrations in the range of 1 pM–10 mM) and apyrase (0.1 U/ml final concentration) were added to give a final volume of 220 μL. Non-specific binding was determined by addition of alprenolol to the negative control (100 μM final concentration). Samples were incubated at 20°C for 1 h, before adding [^3^H]-DHA (concentrations of the competing ligand were varied depending on the apparent K_D_ determined for the construct with and without G protein (see Table S3), so that concentrations of competing ligand were in the range 1-2.5 x K_D_). Samples were incubated at 20°C for 1-5 h (longer incubation times were required for the wild-type β_2_AR to allow equilibration with DHA), before filtering through 96-well fibre filter plates as previously described. Radioligand binding was quantified by scintillation counting and K_i_ values were determined using GraphPad Prism version 7.0b. All K_i_ values obtained are given in Tables S2 and S3 and are mean values obtained from at least two experiments performed in duplicate. Racemic mixtures of the tested ligands were used in all cases apart from measurements with epinephrine and norepinephrine where the active (*R-*) enantiomers were used, and also dobutamine where the chiral β-OH is not present. Where racemic mixtures were used, ligand concentrations and K_i_ values were not corrected for the presence of the inactive enantiomers

#### Radioligand association experiments

Assays were performed using insect cell membranes containing βAR resuspended in assay buffer (20 mM HEPES pH 7.5, 50 mM NaCl, 2.5 mM MgCl_2_, 0.1% BSA). For β_1_AR, [^3^H]-DHA association time course experiments were performed at 4°C, reactions were initiated by dilution of insect cell membranes into [^3^H]-DHA to give final concentrations of approximately 8-fold K_D_ for DHA (β_1_AR, 10 nM, β_1_AR(F325Y) 8.3 nM, 9.2nM). Aliquots (50 μL) were withdrawn at the indicated times and filtered to separate bound from unbound [^3^H]-DHA using Whatman GF/B filters which were further processed as described previously. Experiments were performed in triplicate, with non-specific binding determined by addition of alprenolol to negative controls (1 mM final concentration). Comparisons of relative accessibilities of the ligand binding pocket (LBP) of different βAR subtypes and mutants in the presence of mini-G_s_ were performed at room temperature with [^125^I]-cyanopindolol (Cyp). Samples of receptor (0.06-0.08 nM in final volume 108 μL with and without mini-G_s_ [27.8μM]) were incubated for at least 2 hours at room temperature before addition of 12 μL [^125^I]-Cyp to final concentrations that varied from 750-980 pM. After allowing 1.25 h for binding of [^125^I]-Cyp, 50 μL aliquots were withdrawn and bound [^125^I]-Cyp was separated from unbound by filtration with Whatman GF/B filters. Relative accessibility of the LBP was calculated as a percentage of [^125^I]-Cyp binding, mean values were calculated from 6-7 separate experiments for each construct and are displayed in Table S4.

**Fig. S1.**
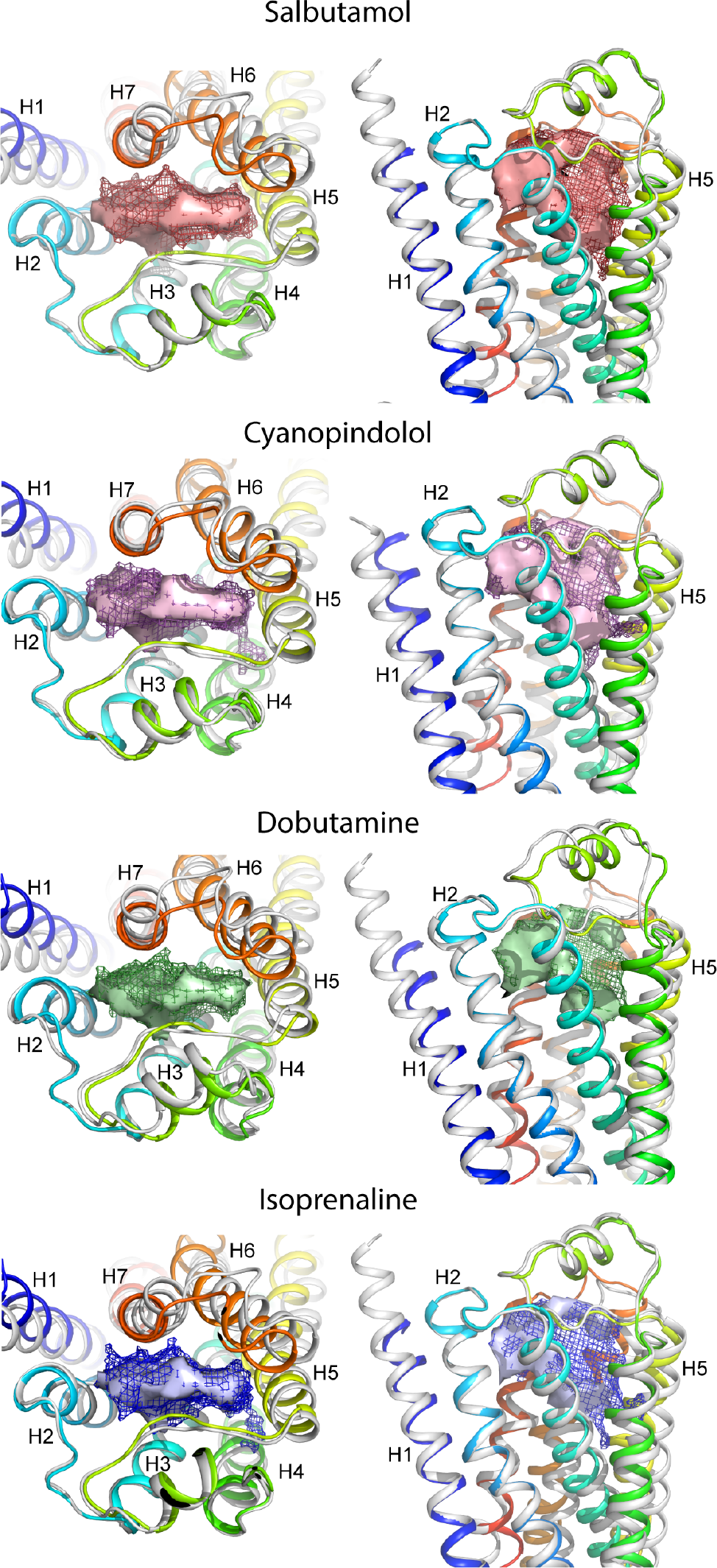
Volume differences of the orthosteric binding site between the active state and active state. In each panel β_1_AR is shown as a cartoon (inactive state, grey; active state, rainbow coloration, N-terminus blue, C-terminus red). The volume of the inactive state is outlined as a mesh and the volume of the active state is outlined as a solid surface. Views on the left are from the extracellular surface and the views on the right are parallel to the membrane plane

**Fig. S2.**
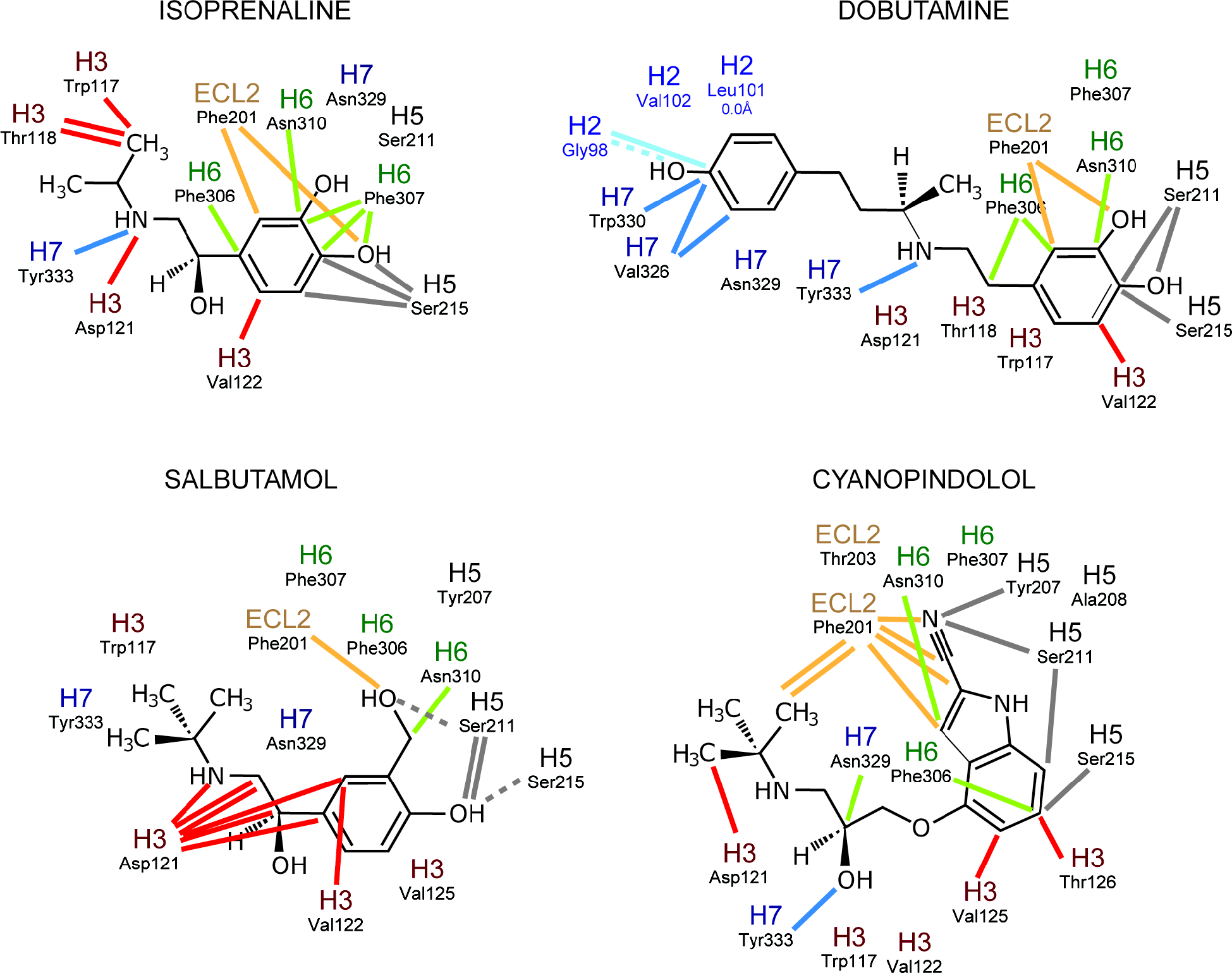
Additional contacts between β_1_AR and ligands in the active state. Structures of agonists co-crystallised with β_1_AR are depicted with receptor-ligand contacts present in the active state, but not in the inactive state, depicted: solid lines, van der Waals interactions (≤ 3.9 Å), dashed line, polar interaction. Colours represent the α-helices or extracellular loops where each residue is located.

**Fig. S3.**
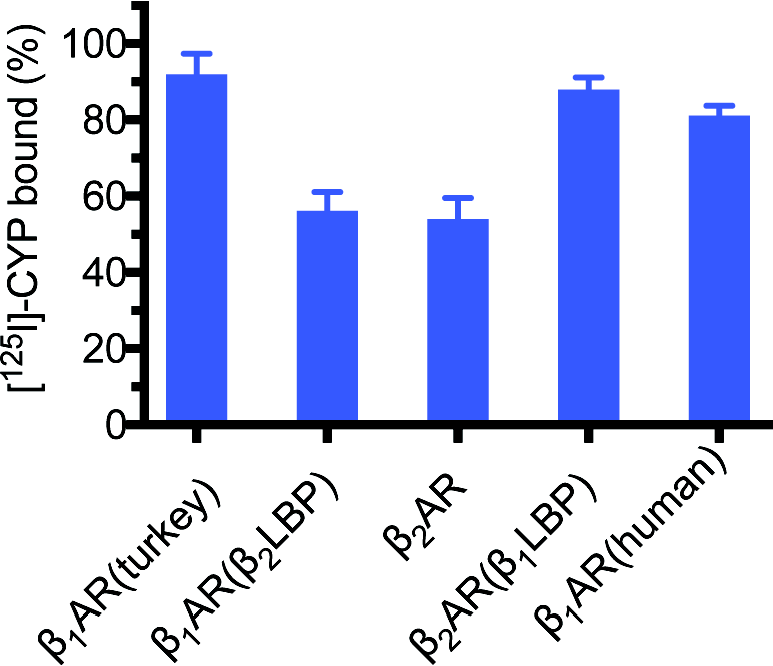
Accessibility of the orthosteric binding pocket to ^125^I-cyanopindolol. The amount of ^125^I-cyanopindolol (^125^I-Cyp) that associated with receptor-mini-G_s_ complexes after a 75 minute incubation (see Methods) was determined in relation to the amount of ^125^I-Cyp bound to the respective receptor in the absence of mini-G_s_. Data represent the mean of 2 independent experiments performed in duplicate.

**Fig. S4.**
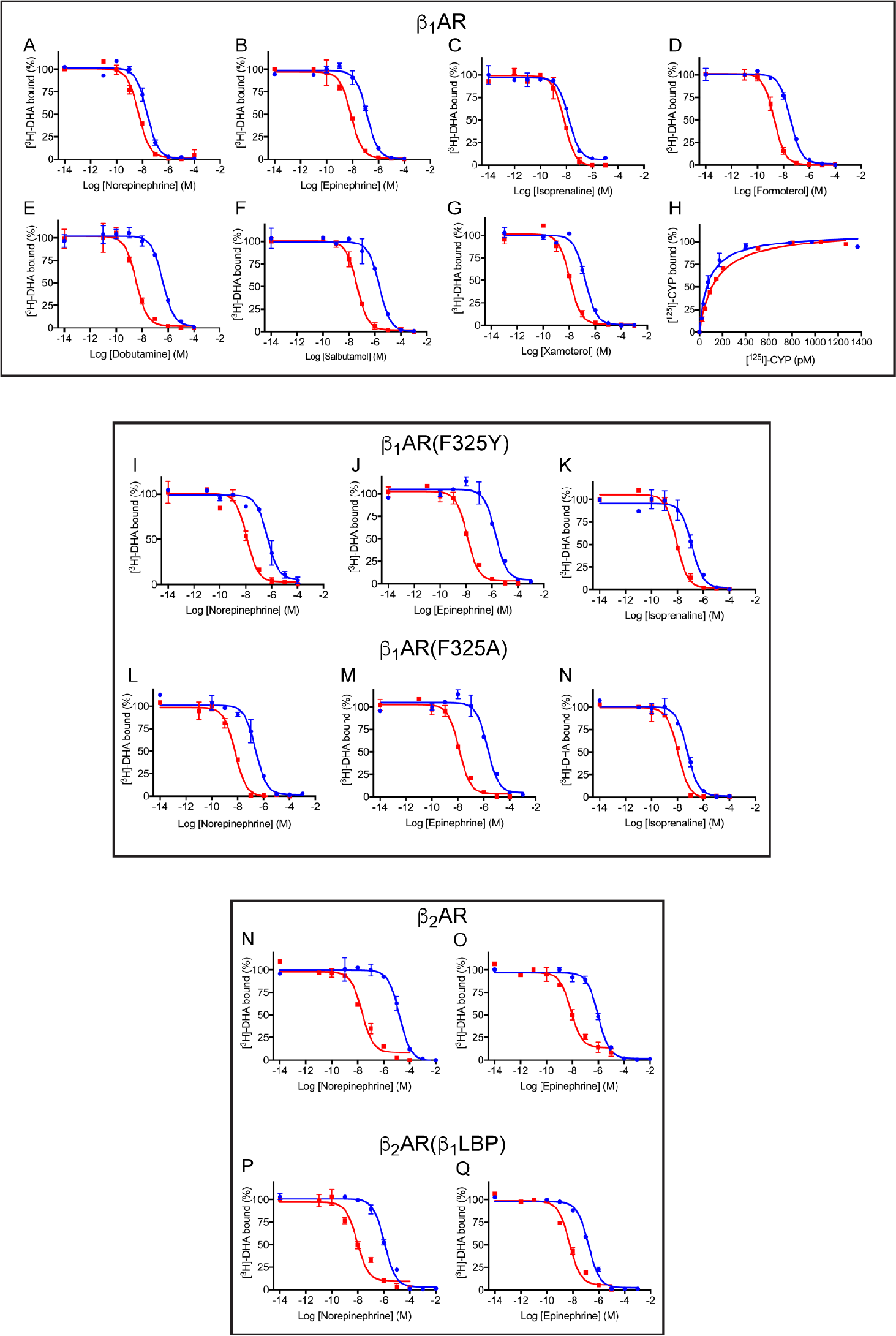
Pharmacology of high affinity and low affinity states. Representative competition binding curves and saturation binding curves are shown for results in Tables S2 and S3. All experiments were performed in duplicate. Experiments to determine the high affinity state were performed in a molar excess of mini-G_s_ (see methods); red curves, low affinity state; blue curves, high affinity state.

**Fig. S5.**
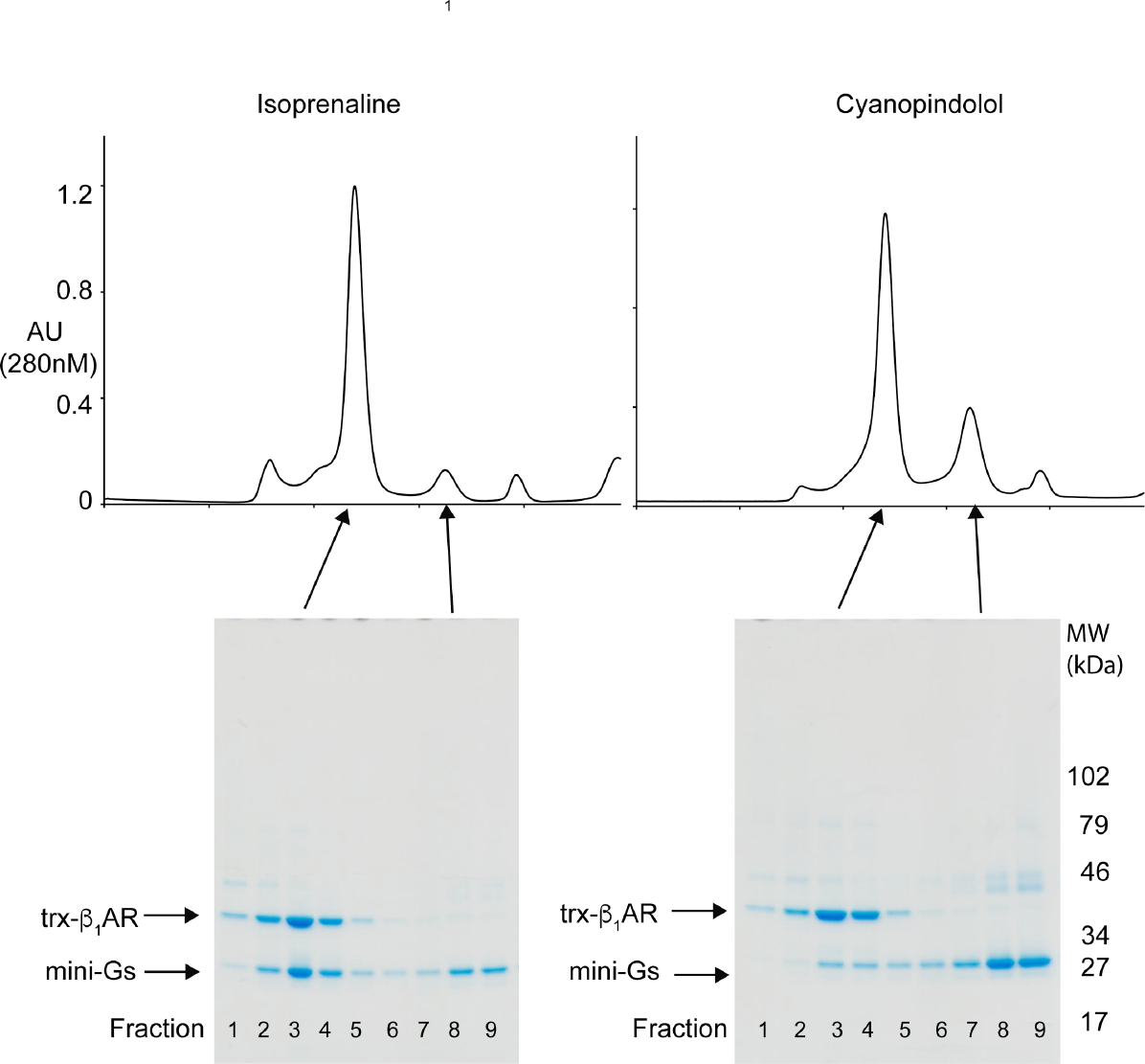
Formation of complexes between trx-β_1_AR and mini-G_s_ in the presence of isoprenaline or cyanopindolol. Complexes were formed as described in the Methods section in the presence of either the full agonist isoprenaline or the weak partial agonist cyanopindolol. The components were then resolved by SEC and the fractions analysed by Coomassie blue-stained SDS-PAGE.

**Fig. S6.**
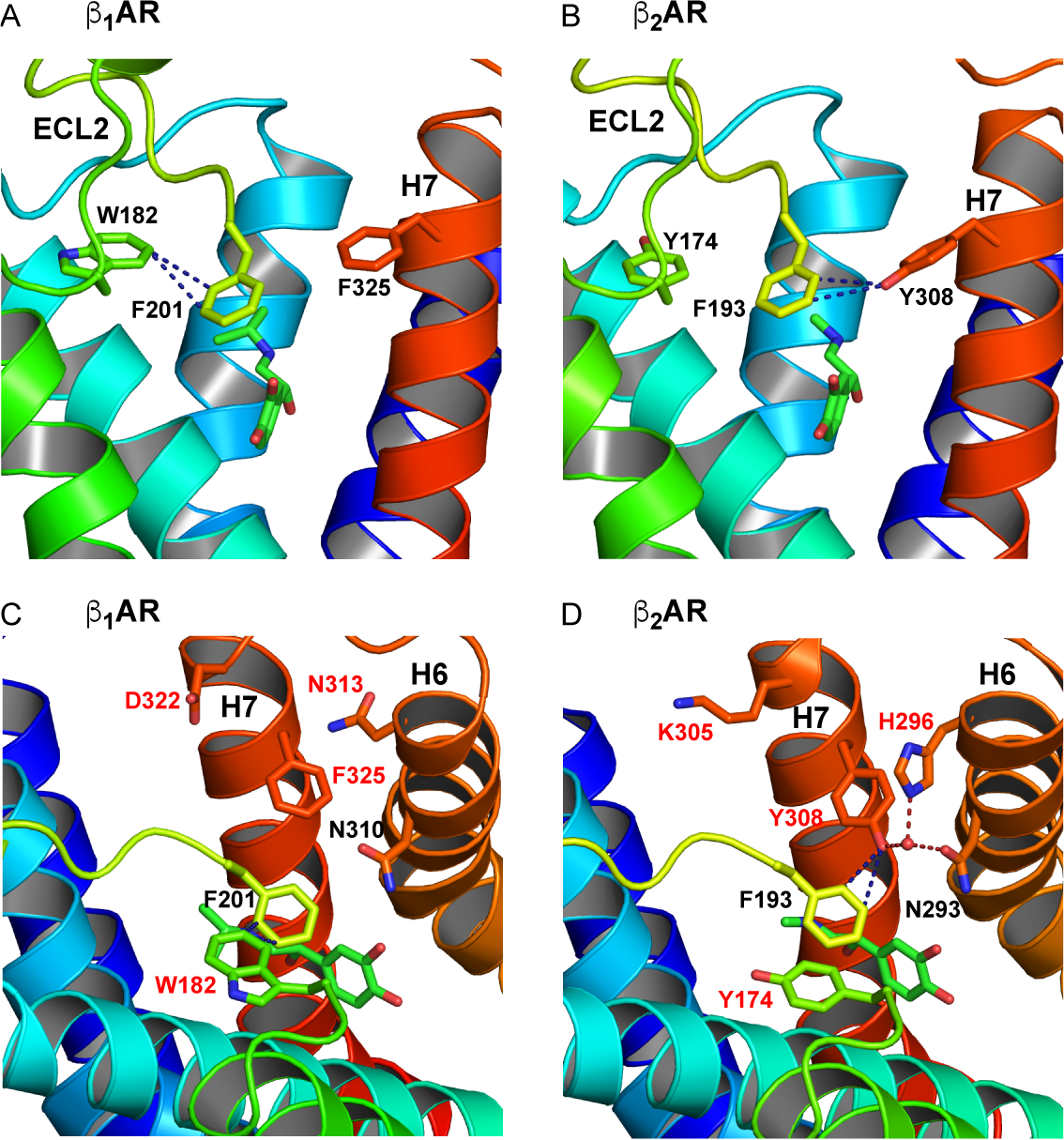
Comparison of β_1_AR and β_2_AR structures: subtype-specific differences imply rationale for mutagenesis. Comparison of structures of (a) activated β_1_AR with isoprenaline bound and (b) β_2_AR with adrenaline bound (4LDO). In β_1_AR, F201 interacts with W182 on ECL2, and not F325 on H7, in the β_2_AR F193 interacts with Y308 on H7, but not with Y174 on ECL2. This subtype specific difference in interactions between F201 (β_1_AR) and F193 (β_2_AR) can be observed in most structures with the exception of β_2_AR crystallized with ligands with bulky headgroups, where interactions between F193 and Y174 can be observed. Alternative views of activated β_1_AR with isoprenaline bound (c) and (d) β_2_AR with adrenaline bound (4LDO) with ECL2 removed for clarity, residues that differ between the two receptors are labeled in red. In the β_2_AR, all of these have been suggested as being involved in high affinity agonist binding states. In the case of H296, this is by participation in an extended H-bond network that also includes T195 on ECL2 (not shown).

**Table S1.** X-ray data collection and refinement statistics

| Ligand, nanobody | Isoprenaline,<br>Nb80 | Formoterol,<br>Nb80 | Salbutamol,<br>Nb6B9 | Dobutamine,<br>Nb6B9 | Xamoterol,<br>Nb6B9 | Cyanopindolol,<br>Nb6B9 |
| --- | --- | --- | --- | --- | --- | --- |
| **ESRF beamlines** | id23-2 | MASSIF-1 | id30b | id30-a3 &<br>id29 | id23-2 | id30-a3 |
| **Number of crystals** | 1 | 1 | 2 | 4 | 1 | 2 |
| **Space group** | P 2<sub>1</sub> 2<sub>1</sub> 2<sub>1</sub> | P 2<sub>1</sub> 2<sub>1</sub> 2<sub>1</sub> | P 2<sub>1</sub> 2<sub>1</sub> 2<sub>1</sub> | P 2<sub>1</sub> 2<sub>1</sub> 2<sub>1</sub> | P 2<sub>1</sub> 2<sub>1</sub> 2<sub>1</sub> | P 2<sub>1</sub> 2<sub>1</sub> 2<sub>1</sub> |
| **Cell dimensions *a*,**<br>***b*, *c* (Å)** | 116.7, 121.2,<br>129.5 | 116.6, 121.1,<br>129.8 | 116.6, 121.5,<br>130.4 | 116.5, 119.7,<br>129.2 | 116.4, 121.5,<br>130.2 | 116.6, 120.0,<br>130.1 |
| **Resolution range** | 41.38-2.78<br>(2.88- 2.78)<sup>1</sup> | 44.28-2.7<br>(2.79-2.7) | 25.77-2.61<br>(2.68-2.61) | 41.08-2.7<br>(2.79-2.7) | 41.5 -2.5<br>(2.56-2.5) | 38.22-2.79<br>(2.89-2.79) |
| **Unique reflections** | 46114 (3946) | 50611 (4416) | 56733 (4129) | 50091 (4298) | 64594 (4470) | 45172 (3688) |
| **Completeness**<br>**before truncation**<br>**(%)** | 98.5 (87.2) | 99.0 (95.3) | 99.0 (89.3) | 99.4 (94.1) | 100.0 (99.7) | 98.3 (82.9) |
| **Multiplicity** | 9.1 (6.4) | 4.6 (4.7) | 9.5 (6.2) | 15.8 (8.6) | 11.9 (11.8) | 9.8 (3.7) |
| **Mean I/σI** | 5.1 (0.0) | 6.6 (1.7) | 5.0 (0.3) | 4.9 (0.5) | 5.4 (0.4) | 3.7 (0.2) |
| **R-merge** | 0.408 (-71.6) | 0.151 (0.936) | 0.334 (6.69) | 0.553 (6.61) | 0.597 (10.3) | 0.378 (5.28) |
| **Resolution limits**<br>**CC1/2=0.3 h, k, l**<br>**axes & overall (Å)** | 2.78, 3.72,<br>3.24, **3.06** | 2.7, 3.62,<br>3.43, **2.92** | 2.67, 3.47,<br>3.42, **3.01** | 2.7, 3.99,<br>3.44, **3.06** | 2.54, 3.83,<br>3.21, **2.99** | 2.91, 4.24,<br>3.63, **3.2** |
| **REFINEMENT** |  |  |  |  |  |  |
| **Resolution (Å)** | 41.4-2.8<br>(2.873-2.8) | 88.6-2.7<br>(2.78-2.7) | 25.8-2.76<br>(2.83-2.76) | 41.1-2.7<br>(2.77-2.7) | 41.5-2.5<br>(2.565-2.5) | 38.2-2.8 (2.87<br>- 2.8) |
| **Completeness,**<br>**truncated data (%)** | 64.88 (9.03) | 62.71 (2.78) | 66.76 (3.85) | 58.75 (12.61) | 56.1 (11.4) | 52.1 (2.94) |
| **No. reflections** | 28351 | 30378 | 30724 | 28040 | 34374 | 23793 (162) |
| **R-work/R-free (%)** | 0.284/0.317<br>(0.333/0.392) | 0.242/0.276<br>(0.323/0.348) | 0.265/0.285<br>(0.427/0.324) | 0.241/0.278<br>(0.452/0.412) | 0.248/0.266<br>(0.418/0.425) | 0.240/0.274<br>(0.496/0.703) |
| **No. atoms** | 8094 | 8271 | 8231 | 8280 | 8211 | 8190 |
| **Protein** | 7934 | 7984 | 8015 | 8014 | 8016 | 7977 |
| **Ligands &**<br>**detergents** | 140 | 258 | 203 | 252 | 170 | 206 |
| **Water** | 20 | 29 | 13 | 14 | 25 | 7 |
| **B-factors (Å<sup>2</sup>)** |  |  |  |  |  |  |
| **Protein** | 73.7 | 70.2 | 84.4 | 64.6 | 62.1 | 83.1 |
| **Ligand &**<br>**detergents** | 54.6, 54.4 | 57.9, 73.8 | 64.1, 77.1 | 57.4, 67.6 | 49.1, 68.6 | 56.5, 78.1 |
| **Waters** | 38.2 | 38.5 | 41.2 | 23.6 | 36.3 | 38.9 |
| **R.M.S.D.** |  |  |  |  |  |  |
| **Bond lengths**<br>**(Å)** | 0.008 | 0.008 | 0.008 | 0.008 | 0.008 | 0.008 |
| **Bond angles**<br>**(°)** | 1.12 | 1.17 | 1.2 | 1.18 | 1.12 | 1.11 |

**Table S2.** Affinities (K_i_s) of β_1_AR and β_2_AR constructs for ligands

| Receptor and construct | Ligand | $K_i$ low affinity state (Log) | SEM | n | $K_i$ high affinity state (Log) | SEM | n | Affinity shift (Log) | Affinity shift (fold) |
| --- | --- | --- | --- | --- | --- | --- | --- | --- | --- |
| $\beta_1$AR | Isoprenaline | -8.16 | 0.01 | 2 | -8.62 | 0.02 | 2 | 0.46 | 2.9 |
| $\beta_1$AR(F325Y) | Isoprenaline | -7.28 | 0.05 | 2 | -8.69 | 0.01 | 2 | 1.27 | 18.7 |
| $\beta_1$AR(F325A) | Isoprenaline | -7.68 | 0.02 | 2 | -8.59 | 0.08 | 2 | 0.9 | 8 |
| $\beta_1$AR | Formoterol | -7.99 | 0.09 | 3 | -9.31 | 0.04 | 3 | 1.32 | 22 |
| $\beta_1$AR | Salbutamol | -5.92 | 0.06 | 3 | -7.80 | 0.06 | 3 | 1.88 | 76 |
| $\beta_1$AR | Dobutamine | -6.86 | 0.03 | 4 | -8.97 | 0.03 | 4 | 2.10 | 126 |
| $\beta_1$AR | Xamoterol | -7.07 | 0.05 | 2 | -8.46 | 0.01 | 2 | 1.38 | 24 |
| $\beta_1$AR | Norepinephrine | -8.09 | 0.06 | 4 | -8.91 | 0.07 | 4 | 0.82 | 6.6 |
| $\beta_1$AR(F325Y) | Norepinephrine | -6.92 | 0.09 | 6 | -8.49 | 0.06 | 4 | 1.58 | 38 |
| $\beta_1$AR(F325A) | Norepinephrine | -7.09 | 0.05 | 2 | -8.76 | 0.01 | 2 | 1.67 | 46.4 |
| $\beta_1$AR | Epinephrine | -7.25 | 0.06 | 5 | -8.51 | 0.03 | 6 | 1.27 | 18 |
| $\beta_1$AR(F325Y) | Epinephrine | -6.20 | 0.06 | 4 | -8.48 | 0.07 | 4 | 2.28 | 189 |
| $\beta_1$AR(F325A) | Epinephrine | -6.56 | 0.05 | 2 | -8.45 | 0.03 | 2 | 1.89 | 77.8 |
| $\beta_2$AR | Norepinephrine | -5.38 | 0.05 | 4 | -8.29 | 0.07 | 7 | 2.91 | 818 |
| $\beta_2$AR($\beta_1$LBP) | Norepinephrine | -6.48 | 0.07 | 3 | -8.57 | 0.05 | 5 | 2.09 | 124 |
| $\beta_2$AR | Epinephrine | -6.47 | 0.03 | 6 | -8.69 | 0.02 | 5 | 2.22 | 166 |
| $\beta_2$AR($\beta_1$LBP) | Epinephrine | -7.31 | 0.01 | 2 | -8.76 | 0.05 | 2 | 1.45 | 28 |

**Table S3.** Apparent K_D_s of β_1_AR and β_2_AR constructs for cyanopindolol and dihydroalprenolol

| Receptor and construct | Ligand | $K_D$ control (Log) | SEM | n | $K_D$ + mini-$G_s$ (Log) | SEM | n | Affinity shift (Log) | Affinity shift (fold) |
| --- | --- | --- | --- | --- | --- | --- | --- | --- | --- |
| $\beta_1$AR | [$^{125}$I]-cyanopindolol | -9.99 | 0.03 | 4 | -9.85 | 0.02 | 4 | -0.14 | -0.72 |
| $\beta_1$AR | [$^3$H]-DHA | -8.87 | 0.08 | 5 | -9.47 | 0.05 | 5 | n/a | n/a |
| $\beta_1$AR(F325Y) | [$^3$H]-DHA | -9.03 | 0.01 | 2 | -9.60 | 0.05 | 2 | n/a | n/a |
| $\beta_1$AR(F325A) | [$^3$H]-DHA | -8.96 | 0.01 | 2 | -9.77 | 0.04 | 5 | n/a | n/a |
| $\beta_2$AR | [$^3$H]-DHA | -9.66 | 0.09 | 3 | -10.19 | 0.03 | 3 | n/a | n/a |
| $\beta_2$AR($\beta_1$LBP) | [$^3$H]-DHA | -9.48 | 0.09 | 4 | -9.88 | 0.03 | 2 | n/a | n/a |

**Table S4.**
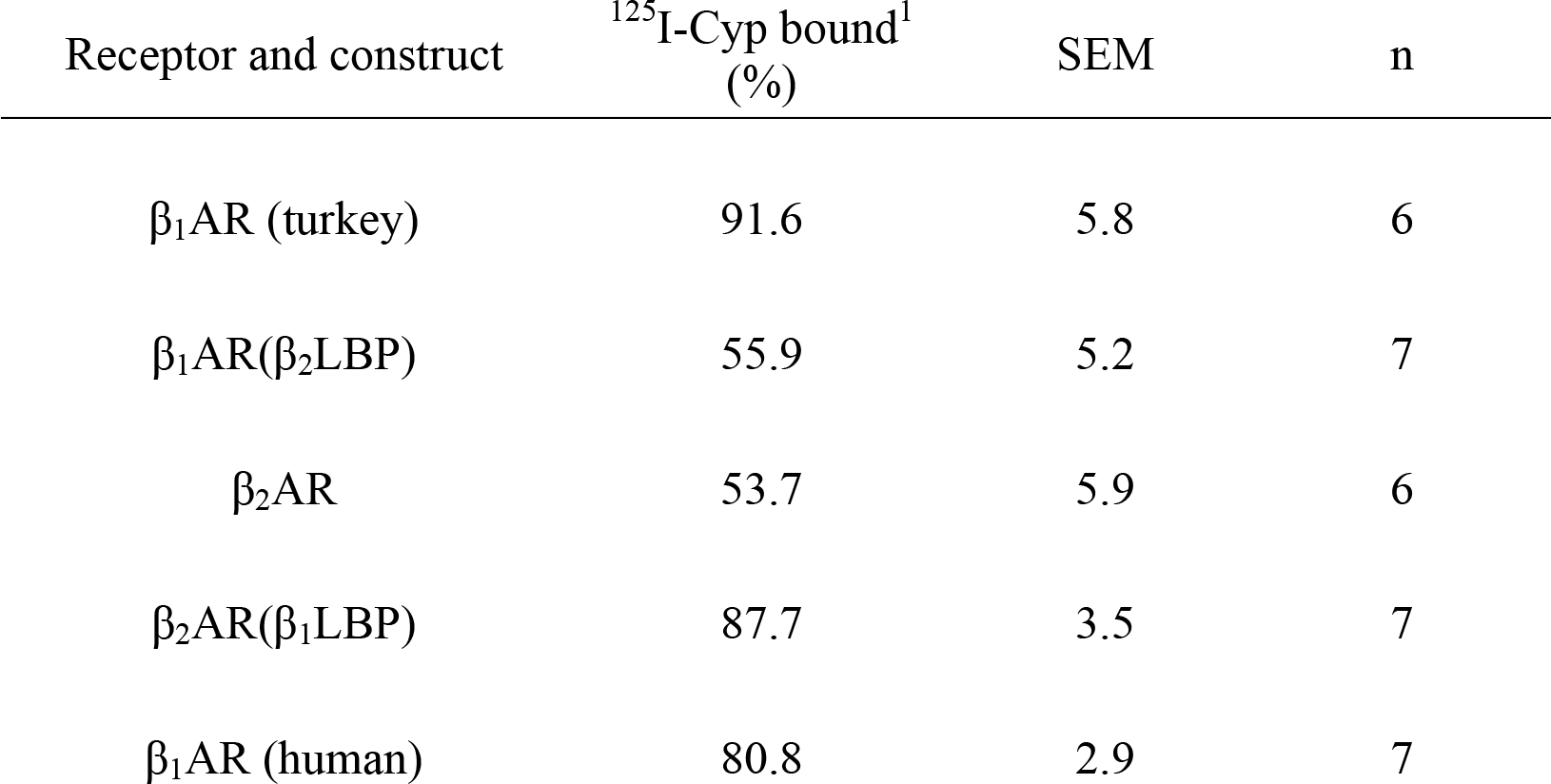
Association of ^125^I-Cyp to active state receptors

**Other Supplementary Materials for this manuscript include the following:**

Data S1 Atomic contacts between receptor and ligands

